# Photoswitchable binders enable temporal dissection of endogenous protein function

**DOI:** 10.1101/2023.09.14.557687

**Authors:** Michael Westberg, Daesun Song, Vandon Duong, Daniel Fernandez, Po-Ssu Huang, Michael Z. Lin

## Abstract

General methods for spatiotemporal control of specific endogenous proteins would be broadly useful for probing protein function in living cells. Synthetic protein binders that bind and inhibit endogenous protein targets can be obtained from nanobodies, designed ankyrin repeat proteins (DARPins), and other small protein scaffolds, but generalizable methods to control their binding activity are lacking. Here, we report robust single-chain photoswitchable DARPins (psDARPins) for bidirectional optical control of endogenous proteins. We created topological variants of the DARPin scaffold by computer-aided design so fusion of photodissociable dimeric Dronpa (pdDronpa) results in occlusion of target binding at baseline. Cyan light induces pdDronpa dissociation to expose the binding surface (paratope), while violet light restores pdDronpa dimerization and paratope caging. Since the DARPin redesign leaves the paratope intact, the approach was easily applied to existing DARPins for GFP, ERK, and Ras, as demonstrated by relocalizing GFP-family proteins and inhibiting endogenous ERK and Ras with optical control. Finally, a Ras-targeted psDARPin was used to determine that, following EGF-activation of EGFR, Ras is required for sustained EGFR to ERK signaling. In summary, psDARPins provide a generalizable strategy for precise spatiotemporal dissection of endogenous protein function.

## Main

A remaining challenge in biology is the direct control of unmodified natural proteins with spatiotemporal precision. Thus, tools for fine control of where and when endogenous proteins are active would have widespread utility in interrogating the roles of specific proteins in biological responses. While genetic tools exist for manipulating endogenous protein abundance over long time courses via gene editing or modulation of mRNA levels^1,2^; generalizable methods for rapidly controlling endogenous protein activity are lacking. Small molecule inhibitors can provide a degree of temporal control, but they are unavailable for many targets and their effects cannot be restricted to genetically defined cells.

Synthetic binding domains (SBDs) are compact protein folds that have been developed as antibody alternatives for efficient recognition of both extra- and intracellular target proteins^3^. SBDs can be rapidly engineered to bind a target by selection^4–7^. Specifically, a defined paratope region of the SBD is diversified to bind targets while the rest of the scaffold is kept constant to preserve the fold. High-affinity SBDs are versatile tools for investigating cell signaling, constructing diagnostic probes, and developing therapeutic biologics^8–15^. By binding to a functionally important site of a target, SBDs can directly inhibit biochemical signaling^14,15^. However, when expressed, the binding activity of SBDs is permanently active. This limits the ability to study dynamic signaling mechanisms. Thus, acute and reversible activation of SBD binding could provide new opportunities for precision perturbation of endogenous targets to study their function.^16^

Photoinducible SBDs that only bind to endogenous targets upon illumination would enable genetically-encodable and acute perturbation of protein function. Examples of photoinducible SBDs have been reported^17– 20^ but are limited in either regulatability or generalizability. Photoreconstitution of SBDs from split fragments can provide robust control under controlled conditions^17^, but basal and induced concentrations of reconstituted SBDs will depend on fragment concentrations, which may vary from cell to cell, and the approach showed negligible binding reversibility. Single-chain designs reduce concentration effects, and have been created by inserting LOV or PYP photodomains into nanobodies, monobodies, and affibodies for allosteric modulation of binding activity^18–20^. However, their utility is hampered by lack of a generalizable photodomain insertion site with predicable effects on binding^18,19^, variable levels of baseline binding and light-inducibility across targets^18– 20^, or the need for exogenous chromophore supply.^20,21^ Finally, while the reported tools can in principle regulate endogenous proteins, few endogenous targets have so far been studied using SBDs with photoinducible binding.^17^

Here we report the development of single-chain photoswitchable DARPins (psDARPins) that are inactive at baseline, are activated by cyan light, and can be inactivated again by violet light. In contrast to previous tools, psDARPins do not require chromophore cofactors, and operate through paratope occlusion rather than an allosteric mechanism. The psDARPin design is modular and robust, and generalization to new targets simply requires replacing the DARPin paratope region. We show that psDARPins can bidirectionally control the kinase activity of endogenous ERK and Ras proteins with temporal precision. Using this approach, we characterized the role of Ras in sustaining ERK activity following EGF-initiated signaling. We found that Ras activity is required for sustained ERK activity, ruling out a role for Ras-independent signaling pathways in sustaining ERK activity.

## Results

Photodissociable dimeric (pdDronpa) has been used to control a range of proteins^22–28^. In these studies, two pdDronpa domains were attached to flank an interaction site so that pdDronpa dimerization would cage the site at baseline. Upon illumination with cyan (∼500-nm) light, pdDronpa switches from a brightly fluorescent dimeric ON-state to a non-fluorescent monomeric OFF-state, thereby exposing the interaction site for protein or substrate binding. The process is reversible, as pdDronpa can be redimerized by reverse photoswitching with violet (∼400-nm) light or by slow (t_1/2_∼1 h) thermal relaxation^24^, resulting in protein recaging.

We set out to apply this general design strategy to DARPins (**Fig. 1a**). DARPins were selected since they are stable, small, and disulfide-free α-helical repeat proteins that contain a single rigid and central paratope and fold robustly in fusion proteins^14,15,29^. They can be selected to bind and inactivate protein targets of interest, and function in both intracellular and extracellular environments. Importantly, the concave paratope is flanked by constant N- and C-terminal cap regions for fusion of the photoswitchable domains. Finally, many DARPins have been reported against a diverse array of targets, and several DARPin-based therapies have entered clinical trials^14,15,29^. Thus, we expected that the successful discovery of a photoswitchable DARPins design would enable optical control over a variety of endogenous proteins.

**Fig. 1.**
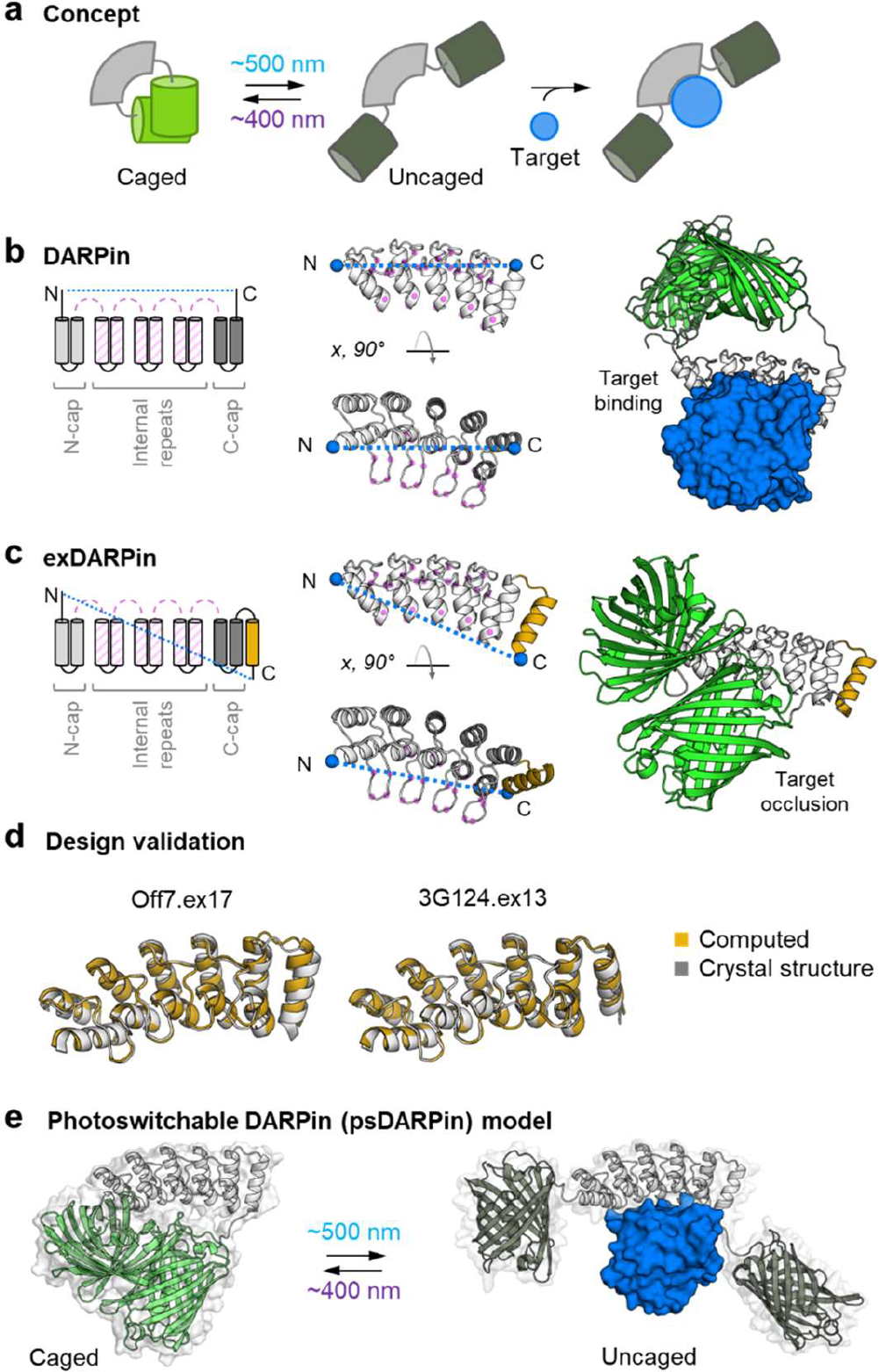
Concept for a generalizable photoinducible synthetic binding protein. **a**, Single-chain optogenetic tools with two brightly fluorescent pdDronpa domains attached to flank a central protein domain allow pdDronpa dimerization and caging of an interaction site at baseline. Photoswitching with cyan light (∼500 nm) leads to loss of pdDronpa fluorescence and uncaging to allow substrate binding. The process is reversible with violet light (∼400 nm). This general concept for photocontrol of binding interactions can be generalized to various targets by fusing pdDronpa to synthetic binding domains (SBDs). **b**, DARPins are cysteine-free SBDs with a centrally located concave binding surface (paratope) that has been selected for high-affinity binding to a variety of protein targets. DARPins are α-helical repeat proteins consisting of at least 1, and typically 2-3, internal repeat units, flanked by N- and C-terminal capping repeats (N- and C-cap). The DARPin paratope is defined by diversifying select residues within the loops and the concave face (magenta spheres). However, fusing pdDronpa to the termini (blue spheres) of the canonical DARPin scaffold is not expected to effectively occlude the paratope. **c**, An extended DARPin (exDARPin) with an additional α-helix appended (gold) to the C-cap is proposed to create a new C-terminal attachment point so pdDronpa dimerization will occur in front of the paratope, effectively caging the exDARPin. **d**, Examples of two RosettaRemodel-designed exDARPins with extensions, ex13 and ex17. Gold is the RosettaRemodel-predicted structure while gray is the experimentally determined crystal structure. Note, the added third C-cap α-helix is packed against the preceding C-cap helices at different angles in the two designs with the C-terminus of ex17 closer to the paratope. **e**, Model of how the photoswitchable DARPin (psDARPin) architecture can bind and inhibit endogenous protein targets upon illumination with cyan light.

### A redesigned DARPin topology

Canonical DARPins are repeat proteins consisting of at least 1, and typically 2-3, internal repeat units^15^ containing the primary paratope, plus N- and C-cap (amino- and carboxy-terminal cap) units that seal and stabilize the scaffold by burying the hydrophobic edges of the internal repeat units (**Fig. 1b**)^15,30,31^. In this canonical DARPin topology, the N- and C-terminal helixes point in the same direction, but this geometry cannot support paratope caging by fusion to pdDronpa (**Fig. 1b**). To enable target occlusion, we redesigned the DARPin scaffold by extending the C-terminal cap with an additional α-helix thus creating a topology with cross-paratope placement of the termini allowing pdDronpa dimerization in front of the paratope (**Fig. 1c)**. Importantly, the redesigned segments of the C-cap do not overlap with the diversified residues that generally form the DARPin paratope (**Extended Data Fig. 1**). Thus, the new topology is not expected to affect target recognition. We hypothesized that this extended DARPin (exDARPin) scaffold would allow photoinducible target binding and modular combination with existing and future DARPins.^15^

Novel exDARPins were generated using computer-aided protein design. Starting from the crystal structure of a model DARPin, GFP-targeted 3G124^32^, RosettaRemodel^33^ was used to generate the novel C-cap of the exDARPins with various loop and helix lengths. Following manual curation of the top-scoring outputs from RosettaRemodel, we selected 18 designs for further experimental validation. C-cap unfolding marks the first event of DARPin denaturating^34^. In solution, exDARPins had high thermal stability with T_m_ > 95 °C (**Extended Data Fig. 2**) exceeding the stability of previously optimized C-caps^34,35^. Crystal structures were obtained for 12 of the extensions after inserting the redesigned C-cap in either 3G124 or Off7, a maltose-binding protein (MBP) targeted DARPin^31^. We chose two designs, ex13 and ex17, for further study. These differed in the positioning of the appended C-terminal helix with the ex17 C-terminus angled closer to the paratope (**Fig. 1d**).

Next, we computationally screened fusions of pdDronpa1.2 (pdD1.2) and exDARPin domains to predict optimal linker lengths for paratope caging using RosettaRemodel^36^ (**Fig. 1e**).To define the tested fusions, we will refer to the exDARPins by the version of their C-cap variant, for example, the exDARPin based on DARPin 3G124 combined with ex13 is 3G124.ex13. Finally, when pdD1.2 is fused to both the N- and C-termini of an exDARPin, we will add a ps-prefix to denote photoswitchability, e.g. ps3G124.ex13. After varying the length of each linker between 1-12 residues, the computations indicated that flexible linkers of 4–7 amino acids at each terminus favored maximal coverage of the paratope (**Extended Data Fig. 3**) in ps3G124.ex13 and ps3G124.ex17. These predictions were used to design the constructs for experimental validation of the psDARPin concept.

### Intracellular validation of the psDARPin architecture

To test the intracellular function of the psDARPins, we developed a live-cell nucleus-cytosol partition assay using the cyan fluorescent protein mTurquoise2 (mTurq2) as a model target (**Fig. 2a**). As mTurq2 is derived from GFP^37,38^, we expected the anti-GFP DARPin 3G124^32^ fused to a nuclear export sequence (NES) could bind mTurq2 and sequester it in the cytosol. To establish the assay baseline, we prepared a negative binding control psOff7.ex0-NES, that does not target mTurq2 but that contains pdD1.2 domains, and a positive binding control 3G124.ex0-NES. Here ex0 refers to a canonical C-cap (**Extended Data Fig. 1a**). When expressed in HeLa cells, 3G124.ex0-NES recruited mTurq2 to the cytosol, while mTurq2 was homogenously distributed in the cells expressing psOff7.ex0 (**Fig. 2b**). These controls demonstrate that mTurq2 recruitment is specific to the 3G124 DARPin, and establish cytosol-to-nuclear (C/N) fluorescence ratios of ∼4.8 for full mTurq2 sequestration and ∼1.0 for no mTurq2 binding (**Fig. 2c**).

**Fig. 2.**
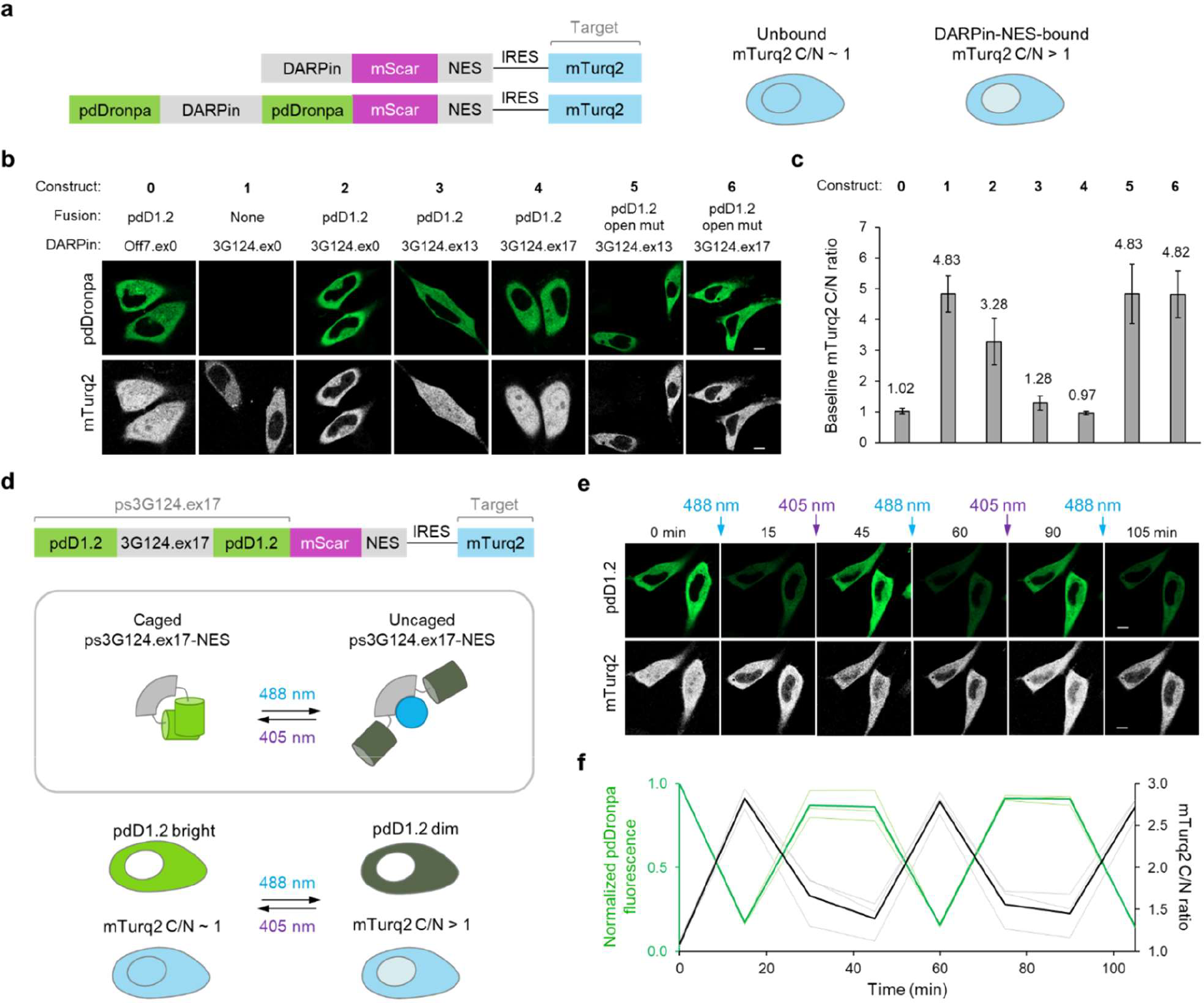
Exploring the caging of extended DARPins. **a**, Domain organization of constructs used for testing DARPin caging by combining DARPin extensions and pdDronpa. DARPins were localized in the cytosol by fusion of a nuclear export sequence (NES), and freely diffusing mTurquoise2 (mTurq2) was expressed as a target of the 3G124 DARPin. Cytosolic localization of the DARPin fusions was validated using a mScarlet tag. **b**, Representative images of HeLa cells expressing test constructs. The constructs are: **0**, the extension-free Off7 (Off7.ex0) DARPin recognizing E. coli maltose-binding protein with terminal pdDronpa1.2 (pdD1.2) fusions, as a negative control not expected to bind mTurq2; **1**, the extension-free GFP-binding 3G124 (3G124.ex0) without pdD1.2 attachments, as a DARPin expected to constitutively sequester mTurq2 in the cytosol; **2**, 3G124.ex0 with pdD1.2 attached to both termini, to test the caging ability of a canonical DARPin without C-terminal extensions; **3**, 3G124 with designed extension ex13 and terminal pdD1.2 attachments (ps3G124.ex13), to test the caging improvement; **4**, 3G124 with designed extension ex17 and terminal pdD1.2 attachments (ps3G124.ex17), to test the caging improvement; **5**, the open pdD1.2 N145K I158V mutant of ps3G124.ex13, to test for preserved mTurq2 binding ability of the fusion; and similarly **6**, the open mutant of ps3G124.ex17, to test for preserved mTurq2 binding ability of the fusion. The N- and C-terminus linkers were 7 and 5 amino acids long for all DARPins tested here. **c**, Quantification of the mTurq2 cytosolic/nuclear (C/N) ratio from b, that is positively correlated with mTurq2 binding to the NES-tagged DARPin fusions. Mean C/N ratios are reported (*n,N* = 18 independent cells, 3 independent experiments per construct), and error bars are standard deviation (SD) calculated using *n*. **d**, Top, the NES-tagged photoswitchable mTurq2-binding 3G124.ex17 (ps3G124.ex17) was used to test optical regulation of target engagement. Below, expected relationship between pdD1.2 fluorescence and mTurq2 cytosolic localization. Illumination by 488-nm light switches pdD1.2 into a dim monomeric off-state. The resulting uncaged psDARPin is expected to sequester mTurq2 in the cytosol, increasing the C/N ratio. Subsequent excitation by 405-nm light restores pdD1.2 to a dimeric form emitting bright fluorescence upon 488-nm excitation. Thus, ps3G124.ex17 becomes primed for recaging pending mTurq2 dissociation. Following mTurq2 dissociation and recaging, mTurq2 will diffuse freely throughout the cell, including back into the nucleus. **e**, Example of bidirectional photocontrol of ps3G124.ex17 binding to mTurq2 in HeLa cells. Photoswitching of pdD1.2 to its off-state by 488-nm illumination (10 s, 0.10 mW) was correlated with depletion of mTurq2 from the nucleus, indicating photoinduction of ps3G124.ex17 binding. Confocal illumination at 405 nm (5 s, 0.56 mW) restored pdD1.2 fluorescence, and mTurq2 reaccumulated in the nucleus, indicating recaging of ps3G124.ex17. The reversible process was then repeated. **f**, Quantification of pdDronpa fluorescence and mTurq2 C/N ratios of e, plotted over time for *n = 3* individual cells (thin lines) and their mean (bold line). All scale bars are 10 μm.

Next, we tested the candidate photoswitchable DARPins based on 3G124.ex13 and 3G124.ex17 with computationally suggested linker lengths (**Fig. 2b and Extended Data Fig. 4**) for caging efficiency in the dark. The green pdDronpa fluorescence of NES-fused ps3G124 variants was localized to the cytosol, and mTurq2 fluorescence was homogenously distributed throughout the cell. With mTurq2 C/N values of 0.97–1.47, all the tested linker variants showed paratope caging. In contrast, ps3G124.ex0, which lacks the extended C-cap, was not efficiently caged by pdD1.2, producing a C/N ratio of 3.28 (**Fig. 2c**). Thus, as expected, the exDARPin topology is crucial for supporting paratope caging. For both ps3G124.ex13 and ps3G124.ex17, the 7,5 linker combination performed the best with C/N ratios of 0.97 and 1.28, respectively (**Extended Data Fig. 4**). For subsequent experiments this 7,5 linker combination was used.

Next, to validate that the fusion of pdDronpa did not in itself interfere with mTurq2 binding we fused the monomeric pdD1.2 N145K I158V mutant^24^ to 3G124.ex13 and 3G124.ex17 to open the pdDronpa cage under dark basal conditions. The open state variants of ps3G124.ex13 and ps3G124.ex17 efficiently bound mTurq2 with C/N ratios of ∼4.8 matching the positive binding control.

Finally, we tested if the psDARPins could be bidirectionally and reversibility controlled with light. Focusing on ps3G124.ex17, we tested for bidirectional photocontrol of target binding (**Fig. 2d**). Cyan 488-nm light caused photoswitching of pdD1.2 into its non-fluorescent OFF-state followed by accumulation of mTurq2 in the cytosol indicating successful photoactivation of target binding (**Fig. 2e**). In contrast, violet 405-nm light caused reverse photoswitching of pdD1.2 into its fluorescent ON-state and movement of mTurq2 back into the nucleus. Repeating the procedure saw mTurq2 shuttling back-and-forth from the nucleus and the C/N ratio increasing and decreasing, respectively (**Fig. 2f**). Thus, cyan and violet light reversibly activates and deactivates target binding of the psDARPins.

### Subcellular target recruitment

By anchoring psDARPins at organelle membranes, subcellular accumulation of the target can be performed for proximity induced reactions or to sequester it away from the native site of function. This mechanism of action can be used to regulate DARPin targets when binding itself does not inhibit target function. To test this idea, we fused ps3G124.ex17 to the outer mitochondrial membrane or the plasma membrane. The psDARPins labeled the organelles with the expected localization pattern (**Extended Data Fig. 5**). In the dark, mTurq2 was homogenously distributed in the cells indicating robust caging. Following uncaging with cyan light, mTurq2 was enriched at the location of the psDARPin, while illumination with violet light released mTurq2 to the cytosol (**Extended Data Fig. 5**). These experiments show that a subcellularly anchored psDARPins can mediate reversible bidirectional photocontrol of target localization.

### Regulating endogenous kinase activity

The preceding experiments used mTurq2 as a convenient exogenous target to validate the psDARPin function. However, the major feature of psDARPins would be the ability to directly control endogenous proteins. To demonstrate control of endogenous signaling, we initially focused on ERK1/2 of Ras-Raf-MEK-ERK pathway (**Fig. 3a**, hereafter the ERK and Ras-ERK pathway) that are involved in regulating cell differentiation and proliferation under both normal and pathological conditions^39–42^.

**Fig. 3.**
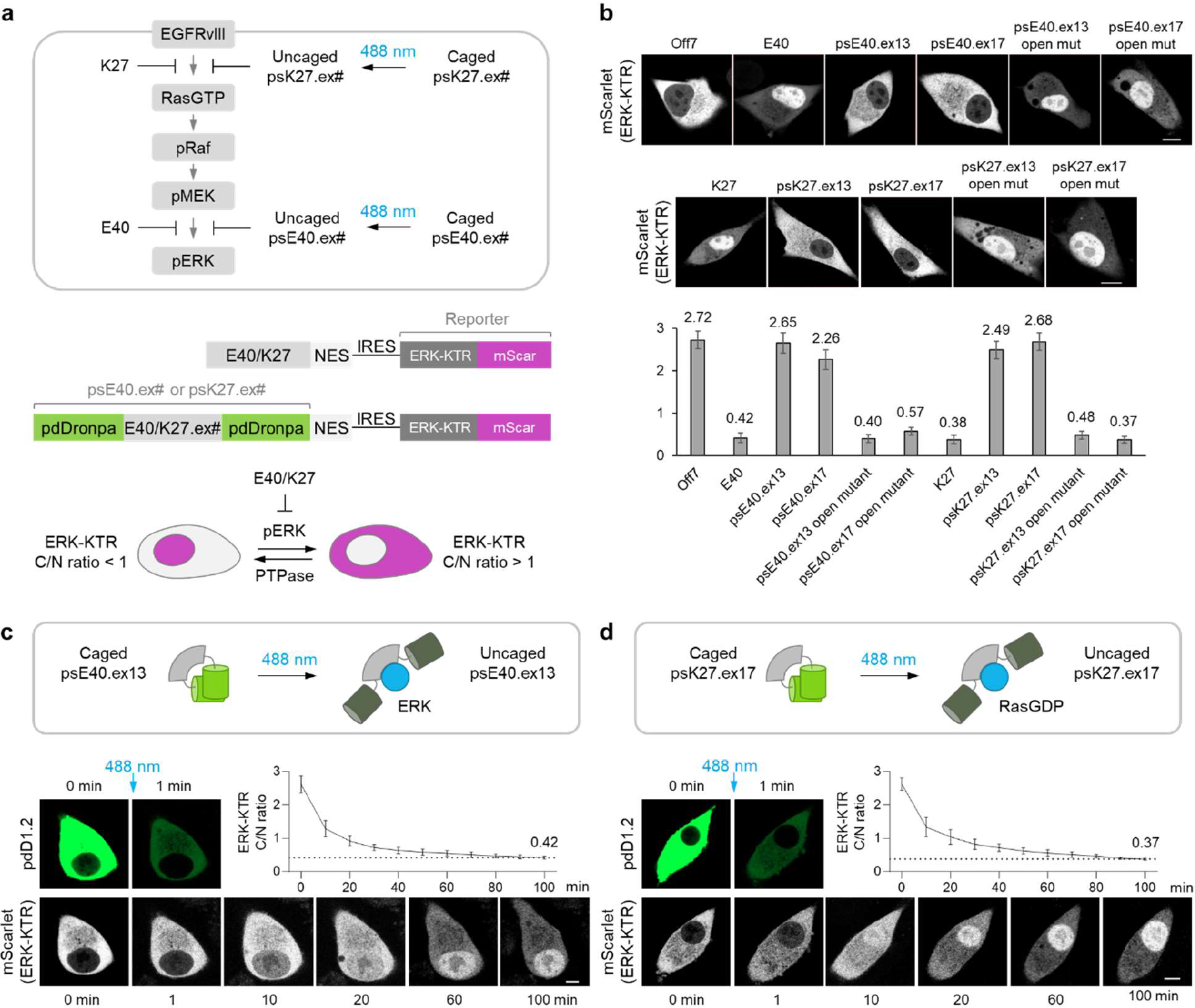
Photoinduced inhibition of endogenous ERK and Ras with psDARPins. **a**, Top, schematic description of Ras-Raf-MEK-ERK signaling pathway in cells expressing the constitutively active EGFRvIII. The enzymes are shown in their active form. The canonical DARPins K27 and E40 are expected to inhibit respectively the formation of active GTP-bound Ras and phosphorylated ERK, RasGTP and pERK, by binding to inactive GDP-bound Ras and unphosphorylated ERK. Middle, NES-tagged psE40.ex# and psK27.ex# were prepared and tested in Ras-ERK hyperactive U87-EGFRvIII cells. Bottom, the activity of ERK and upstream Ras was monitored using a mScarlet-tagged ERK-KTR kinase activity reporter. Active pERK causes ERK-KTR phosphorylation and accumulation in the cytosol, while the reporter is imported into the nucleus following phosphatase-mediated dephosphorylation. Thus, the C/N ratio is positively correlated with ERK activity. **b**, Representative mScarlet images from U87-EGFRvIII cells expressing test constructs under basal dark conditions and quantification of the associated ERK-KTR C/N ratios (*n,N* = 18 independent cells, 3 independent experiments per construct). The negative control Off7 provides a baseline ERK-KTR C/N ratio of ∼2.7 representing full Ras-ERK activity. Expression of E40 caused ERK inhibition and dropped the C/N ratio to ∼0.4. By Comparing to these baselines, the caging ability of psE40.ex13 and psE40.ex17, and the inhibitory efficacy of the respective open (pdD1.2 N145K I158V) mutants thereof were tested. Similarly, K27 was used to establish the baseline for Ras inhibition to allow testing of the caging and binding ability of psK27.ex13 and psK27.ex17 and their open variants, respectively. **c** and **d**, Photoactivation of psE40.ex13 and psK27.ex17 488-nm light (10 s, 0.21 mW) led to gradual inhibition of ERK and Ras with the C/N ratios (*n,N* = 12 independent cells, 3 independent experiments per construct) settling at ∼0.4 (dotted line) similar to the positive inhibition controls, E40 and K27 from panel b. In all instances, scale bars are 10 μm, mean C/N ratios are reported, and error bars are standard deviation (SD) calculated using *n*.

Previous studies showed that the E40 DARPin binds to unphosphorylated ERK^43,44^ and prevents phosphorylation and activation by MEK^43^. However, as E40 had previously been shown only to prevent phosphorylation of chimeric exogenous ERK^43^, we first established that E40 can inhibit endogenous ERK activity in a live-cell assay. As a real-time kinase activity reporter we use mScarlet-labelled ERK-KTR^45^ that moves from the nucleus to the cytosol after phosphorylation by active ERK (i.e., pERK, **Fig. 3a**). Thus, the cytosol-to-nuclear (C/N) mScarlet fluorescence ratio increases with ERK activity^45^. When expressed in Ras-ERK hyperactive human glioma U87-EGFRvIII (U87E) cells together with the non-binding Off7 control, the ERK-KTR was predominantly localized in the cytosol with a C/N ratio of 2.72 (**Fig. 3b**). In contrast, the C/N ratio was 0.42 in E40 expressing cells (**Fig. 3b**), indicating that E40 is indeed capable of inhibiting activation of endogenous ERK.

To design a photoswitchable ERK-inhibiting DARPin, we exchanged the paratope-containing internal repeats of 3G124.ex13 and 3G124.ex17 with the homologous region of E40. We then determined how well the resulting psE40.ex13 and psE40.ex17 were caged in the dark and could be photoinduced. In the dark, psE40.ex13- and psE40.ex17-expressing U87E cells exhibited C/N values of 2.65 and 2.26, respectively (**Fig. 3b**). The C/N ratio for nonilluminated psE40.ex13 is similar to that of Off7 negative control cells, indicating effective caging of the paratope in this design. At 100 min post-illumination with cyan light, the C/N ratio dropped to 0.42 in psE40.ex13-expressing cells (**Fig. 3c**). This is comparable to the C/N value seen in cells expressing constitutively active E40 (**Fig. 3b**) and suggests efficient uncaging of the paratope and preservation of binding activity given enough time for pERK dephosphorylation. Finally, sequential illumination with cyan and violet light confirmed that psE40.ex13 could reversibly control ERK activity (**Extended Data Fig. 6**).

### Inhibiting endogenous wild-type RAS

Ras GTPases are notoriously difficult to inhibit using small molecules^40,46^. With a few exceptions^47–49^, recent progress stems from the discovery of inhibitors targeting specific Ras mutants^40,46^, and a only a few pan-isoform inhibitors of WT Ras have been reported^47,48^. Thus, there is a shortage of specific probes for acute intracellular inhibition of WT Ras. SBDs have larger binding surface areas than small molecule inhibitors, which can facilitate binding to proteins with shallow pockets and flat surfaces. Indeed, several nM affinity monobodies and DARPins have been reported to inhibit WT Ras^50–53^. We focused on DARPin K27 that binds all Ras isoforms (i.e., HRas, KRas, and NRas) in the inactive GDP-bound state and blocks interactions with SOS and RAF thereby inhibiting RAS activation^51,54^.

K27 contains three internal repeat units allowing modular insertion in the validated psDARPin scaffold to create psK27.ex13 and psK27.ex17. Since ERK is activated downstream of Ras, ERK-KTR was also used to monitor Ras activity (**Fig. 3a**). When expressed in U87E cells, K27 established a C/N ratio of 0.38 for constitutively inhibited Ras (**Fig. 3b**). In the dark, psK27.ex13 (C/N 2.49) and psK27.ex17 (C/N 2.68) both match the C/N value of the non-binding Off7 control indicating efficient caging (**Fig. 3b**). Constitutively open variants of psK27.ex13 and psK27.ex17 efficiently bound and inhibited Ras with a C/N ratio similar to K27 (**Fig. 3b**). After uncaging of psK27.ex17 with cyan light, the C/N value decayed to 0.37 at 100 min post-illumination matching the K27 control (**Fig. 3d**). Within the experimental error, the C/N ratio decayed at the same rate following Ras inhibition with psK27.ex17 or ERK inhibition with psE40.ex13. Thus, the rate determining step of this assay in U87E cells is likely either ERK or ERK-KTR dephosphorylation.

### Ras is required for sustained EGFR to ERK signaling

Finally, we used psK27.ex17 to temporally dissect endogenous Ras signaling. Stimulation with EGF causes activation of EGFR at the plasma membrane and a transient burst of Ras-ERK activity. Following the initial burst, endocytosis and deactivation of EGF-bound EGFR and negative feedback in the Ras-ERK signaling pathway leads to a down-regulation of ERK activity for a sustained low level^55–57^, that we refer to as the late phase ERK activity. Various mechanisms have been proposed for late phase signaling from EGFR to ERK^55,56^, however, unequivocal evidence for the role of Ras activity has not been reported. Dynamic signaling of untagged Ras in live cells can be studied by observing Ras activation via biosensors^58–60^ or chemo-/optogenetic activation of Ras^36,61–63^, but acute loss-of-function experiments were, until recently, unfeasible due to the lack of small-molecule inhibitors of wild-type Ras. The unique ability of psK27.ex17 to inactivate endogenous Ras with precise timing allows us to address if Ras activity is required for late phase ERK activity.

We first confirmed that EGFR activation in HeLa cells has a distinct late phase. Addition of 10 ng/mL EGF labelled with Alexa Fluor 647 to serum-starved HeLa cells was followed by internalization within 20 min (**Fig. 4b**), replicating earlier observations^55,56^. We then assessed dynamics of Ras-ERK signaling using the ERK-KTR sensor. Serum-starved HeLa cells have low basal ERK activity (C/N ∼0.5, **Fig. 4c-d**) relative to Ras-ERK hyperactive U87E cells (C/N ∼2.7, **Fig. 3b**). Cells transfected with GFP-binding ps3G124.ex17 were used to establish a control for non-specific effects of psDARPin expression and cyan light illumination. Following stimulation with 10 ng/mL EGF, the C/N ratio saw a transient increase peaking after 30 min (**Fig. 4d**) in accordance with earlier studies using similar conditions^64^. Note; the response of the ERK-KTR reporter is expected to show slower kinetics than pERK levels^56^ and ERK nuclear translocation^65^.

**Fig. 4.**
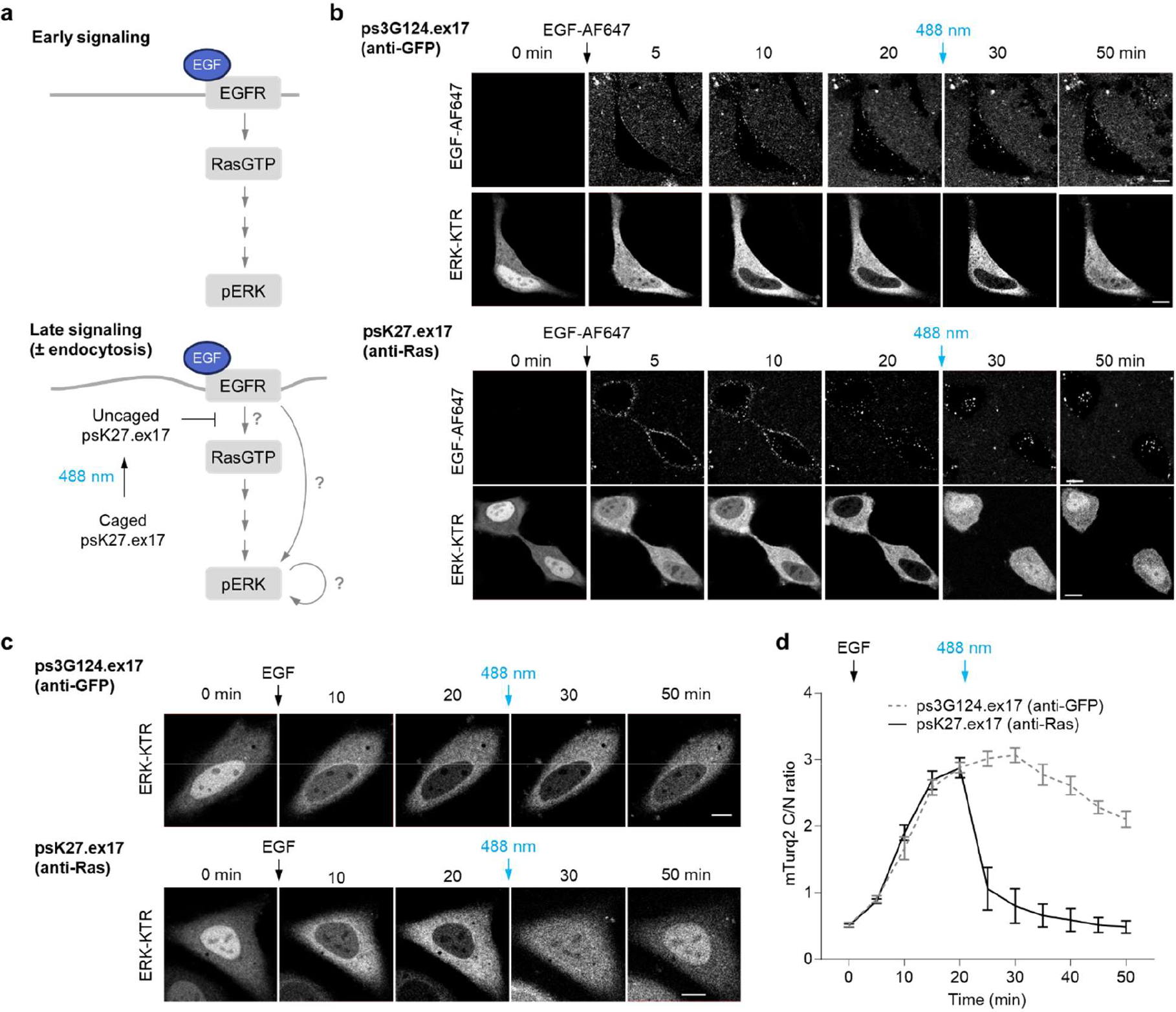
Photoactivated inhibition of endogenous ERK and Ras with psDARPins. **a**, Schematic description of Ras-ERK signaling following stimulation with EGF. In the early burst phase EGF-complexed EGFR activates the Ras-ERK pathway. Subsequently, EGFR endocytosis and negative feedback in the Ras-ERK pathway reduce the ERK activity, but low-level ERK activity is sustained in the late signaling phase. However, the precise role of Ras in late phase signaling is not confirmed. **b**, Representative images of Alexa Fluor 647-labelled EGF (EGF-AF647) used to monitor the dynamics of EGF endocytosis in serum-starved HeLa cells. EGF-AF647 was added at 10 ng/mL to cells co-expressing ERK-KTR and either the negative control ps3G124.ex17 or the Ras-targeted psK27.ex17. Endocytosis was seen within 20 min under both conditions. **c**, Representative images of the ERK-KTR reporter in serum-starved HeLa cells stimulated with 10 ng/mL unlabeled EGF. Immediately after stimulation the reporter beings to move to the cytosol indicating ERK activation with similar behavior under both conditions. Photoswitching with 488-nm light at 20 min post EGF addition immediately caused ERK-KTR to move back into the nucleus in only the psK27.ex17 expressing cells. **d**, Quantification of ERK-KTR C/N ratios from c. In accordance with previous reports, HeLa cells expressing ps3G124.ex17 showed the expected peak in ERK activity at 30 min post EGF stimulation. In contrast, uncaging of psK27.ex17 caused an immediate drop in the ERK activity indicating that Ras activity is required to sustain late phase ERK activity. In b-c, uncaging was performed with 488-nm light (10 s, 0.21 mW). Scale bars are 10 μm, mean C/N ratios are reported (*n,N* = 20 independent cells, 3 independent experiments per construct), and error bars are standard deviation (SD) calculated using *n*.

We next repeated the experiments in cells transfected with Ras-targeted psK27.ex17. Photoactivation of psK27.ex17-transfected cells at 20 min post addition of EGF caused an immediate decay in the C/N ratio that reached baseline level at ∼50 min (**Fig. 4c-d**). These results clearly indicate that Ras activity is required for late phase ERK activity from EGFR to ERK in HeLa cells, dismissing the possibility for alternative mechanisms of EGF-induced signaling to ERK or buffering capacity in the Raf-ERK nodes.

## Discussion

In this study, we developed photoinducible psDARPins for acute control of endogenous proteins. Previously, constitutively active SBDs have been fused to translocation-based optogenetic tools to control endogenous protein activity by either 1) removing targets from their site of function or 2) using photoinduced nuclear to cytosol translocation to control target inhibition at the plasma membrane^65^. The first approach only allows inhibition of targets that can be deactivated by sequestering from their native subcellular site of activity. The second approach only applies to membrane anchored targets. More importantly, while the primary perturbation is controlled by binder translocation, binding is active in the basal state and can perturb target function. For example, an anti-Ras tool reduced Ras-ERK activity even in the dark^65^. In contrast, psDARPins caused no reduction in ERK or Ras activity in the dark. By directly controlling the binding event, psDARPins and other photoinducible SBDs^17–20^ ensure normal target distribution and function prior to illumination and can acutely perturb target function at any subcellular site.

Photoinducible psDARPins rely on an occlusion mechanism for predictable off-on photoswitching of target binding. In contrast, photocontrolled nanobodies or monobodies rely on an allosterically mechanism and require empirical screening of LOV domain insertion to determine the switching direction^18,19,66^. For example, only one photoinducible monobody has been reported^19^, while photorepressed monobodies that bind the target in the dark and release it upon blue-light illumination have been easier to discover.^19,66^ We suggest that these challenges originate in the flexible nanobody and monobody paratopes that primarily rely on diversification of flexible loops^12,67^. Since their paratopes can take many tertiary structures and the loops contribute disparately to the binding of various targets, it is currently not possible to predict how LOV domain insertion will affect the allosteric modulation of binding. This reduces generalizability and modularity of these tools. In contrast, the rigid exDARPins provide a robust scaffold for creating predictable photoinducible binders.

In the design of the exDARPins C-caps, we preserved the canonical DARPin paratope positions to ensure compatibility with existing and future DARPins. Using DARPins targeting either a fluorescent protein, ERK, or Ras, we found that both exDARPin version ex13 and ex17 were simple to integrate and preserved target recognition. Thus, it was straightforward to generate psDARPins that target endogenous kinases. ERK-targeted psE40.ex13 is a new tool for directly controlling ERK activity. While some selective ERK inhibitors have been reported, these do not allow for spatial control of ERK activity^68^. Thus, photoinduced ERK inhibition by psE40.ex13 can complement kinase activating optogenetic tools for investigations of spatiotemporal ERK signaling dynamics^24,61,62,69,70^. Similarly, psK27.ex17 provides an unique tool for pan-isoform inhibition of WT Ras, which can currently not be achieved by any other tool with spatiotemporal precision. In addition, the K27 DARPin can be exchanged with isoform specific anti-Ras DARPins^54^ since the large binding surface of DARPins allows specificity tuning by interactions with non-conserved regions. Similarly, psDARPins may be used to conditionally recognize proteins with post-translational modifications^43,71^, a difficult task for small molecule inhibitors.

Besides inhibition, additional functions can be added to the psDARPins by fusion at the peripheral pdDronpa termini. We used this ability to add subcellular localization tags. Thus, ps3G124.ex17 can be used to relocalize GFP-tagged proteins in the vast amount of available GFP knock-in models. Additionally, fusion to catalytic enzyme domains will allow photoinducible control of proximity-induced reactions. This for example includes the creation of light-controlled degradation by fusing an E3 ligase or light-controlled perturbation of the phosphoproteome by fusing a kinase or phosphatase^72^. Alternatively, another SBD (e.g., another DARPin) may be fused to recruit an endogenous enzyme rather than introducing enzyme-psDARPin chimeras, thus making the system rely on both an endogenous target and an endogenous enzyme.

Possible future applications of psDARPins also include photocontrolled therapeutics. For example, psDARPins could serve as photoinducible antigen receptors for cellular therapies (e.g., CAR-T) or psDARPins could serve as advanced light-activated biologics. In both cases, exogenous optical control allows management of efficacy and side-effects by either local activation of binding or by tuning the dose of active uncaged psDARPins.

## Conclusion

We report the development of psDARPins as single-chain photoinducible binders for control of endogenous proteins. To allow caging of the concave DARPin paratope, we created a novel extended DARPin scaffold, exDARPin, using computational design to add an α-helix to the DARPin C-cap. In contrast to canonical DARPins, the termini of exDARPins are placed in a cross-paratope topology allowing pdDronpa fusions to efficiently cage the paratope and occlude target binding under dark basal conditions. A key feature of the exDARPin design is that the canonical DARPin paratope is preserved. Thus, existing DARPins could easily be incorporated to create a variety of psDARPins. Reversible bidirectional light-mediated control of target binding was demonstrated, and psDARPins were able to acutely inhibit endogenous ERK and Ras. Using psDARPins to temporally dissect EGF-initiated Ras-ERK signaling, we found that Ras activity is required for sustaining ERK activity following EGFR endocytosis in HeLa cells. We expect that the modular psDARPin design will be broadly applicable to existing as well as new DARPins. Thus, the psDARPin architecture provides a modular and generalizable optogenetic method to target endogenous proteins and opens for new ways to investigate and control cell signaling with spatiotemporal precision.

## Methods

### Computational protein design

RosettaRemodel^33^ was used to design the exDARPins and screen the linker lengths between the exDARPin and pdDronpa.

### Cloning

Cloning was performed using standard molecular biology techniques to assemble Q5 DNA polymerase (NEB) amplified or synthesized (IDT) DNA fragments with NEB HiFi Assembly (NEB).

### Bacterial expression and purification

DNA sequences encoding the extended caps were ordered as synthetic e-blocks (IDT) and inserted into two model DARPins: 1) 3G124 (anti-GFP) and 2) and Off7 (anti-MBP) and cloned into a pET vector (Addgene plasmid #29666) containing a N-terminal His-Tag. The two DARPin models were used to facilitate downstream protein crystallization. The DARPin expression plasmids were transformed into T7 Express lysY/Iq E. coli cells (NEB). From single colonies, 50 mL cultures of Auto Induction Media (Formedium) were inoculated and grown at 37 °C and 250 rpm. After 24 h, the cells were harvested, and the pellets were frozen. Chemical lysis of the resuspended pellets was performed in 3 mL/g pellet B-PER II (Thermo Scientific) supplemented with 40 U/mL Pierce universal nuclease (Thermo Scientific) and cOmplete™ EDTA-free Protease Inhibitor Cocktail (Roche). The supernatant was cleared by centrifugation at 12000 × g for 15 min. The soluble fraction was batch-absorbed onto Ni-NTA resin (Thermo Scientific) by adding buffer to the supernatant for a final buffer composition of approximately 40 mM Tris (pH 8.0), 25 mM imidazole, and 500 mM NaCl. The resin was loaded onto gravity flow columns and washed with 20 column volumes of a buffer containing 50 mM Tris (pH 8.0), 25 mM imidazole, and 500 mM NaCl. High purity protein was eluted in a buffer of 50 mM Tris (pH 8.0), 250 mM imidazole, 500 mM NaCl, and 10 mM EDTA. Eluted fractions with high protein content were pooled and buffer exchanged into TBS (20 mM Tris pH 7.4, 136 mM NaCl) using Econo-Pac 10DG desalting columns (Bio-Rad). Purity of the samples was checked on SDS-PAGE (>95%), and protein concentrations were determined based on A280 and predicted extinction coefficients.

### Crystallography

3G124.ex# and Off.ex# DARPins were concentrated to 10 mg/mL in TBS using Amicon Ultra-4 10K centrifugal filters (Millipore). Sitting drops were dispensed on Intelli-plates 96-3 LVR (Hampton Research) using an Oryx8 crystallization robot (Douglas Instruments). The plates were incubated at 6 °C and examined using RockImager2 UV-Vis Imager (Formulatrix). Crystals formed in various wells of the commercial Morpheus HT-96 screen. Crystals were harvested and added to precipitant solution mixed with cryoprotectant and cryocooled by plunging into liquid N2.

Data collections were performed at 100 K using Stanford Synchrotron Radiation Lightsource (SSRL) beamlines BL12-1 (SLAC National Accelerator Laboratory, Menlo Park, USA) (5). Data were reduced using XDS^73^ or DIALS^74^, scaled using AIMLESS^75^, and analyzed with different computing modules within the CCP4 suite^76^ or CCP4i2 suite^77^. Crystals belonged to the monoclinic P1211 or I121 space groups and contained one polypeptide chain per asymmetry unit. Structures were solved by the molecular replacement method with Phaser^78^ using the RosettaRemodel outputs as the search model. Refinement was performed using REFMAC^79^ and manual refinement performed in COOT^80^.

### Differential scanning fluorimetry

The thermal stability of 3G124.ex# DARPins was tested using differential scanning fluorimetry. In white bottom 96 well plates (Bio-Rad), 50x concentrate SYPRO Orange dye (Invitrogen) was added to 10 μM protein in TBS. In a CFX96 Touch Real-Time PCR instrument (Bio-Rad), 50 uL samples were heated from 20 to 100 °C at 0.5 °C/min while the fluorescence of SYPRO Orange was monitored in the FRET channel. Fluorescence data were analyzed in MATLAB (Mathworks). Background subtraction was performed using the data from a well only containing SYPRO Orange in TBS. The thermal melting temperature was determined from the position of the maximum of the first derivative of the fluorescence signal.

### Mammalian cell culture and transfection

HeLa cells (ATCC, CCL-2) were cultured at 37 °C under a 5% CO_2_ humidified atmosphere in Dulbecco’s Modified Eagle’s Medium (DMEM, Gibco), supplemented with 10% fetal bovine serum (FBS, Gemini), 2 mM L-glutamine (Gibco), 100 U/ml penicillin, and 100 μg/ml streptomycin (Gemini). U87-EGFRvIII (U87E) cells, that express the constitutively active EGFR variant III^81^, were previously developed in the Lin lab by pcDNA3-EGFRvIII transfection of U87 cells and Geneticin selection. U87E cells were maintained in DMEM supplemented with 10% bovine calf serum (BCS, Gemini), 2 mM L-glutamine (Gibco), 100 U/ml penicillin, 100 μg/ml streptomycin (Gemini), 0.8 mg/mL Geneticin (Gibco), 10 μg/ml Ciprofloxacin (Sigma), and 10 μg/ml Piperacillin (Sigma) at 37 °C under a 5% CO2 humidified atmosphere. HeLa and U87E cells were seeded into 8-well slide chambers (ibidi) at a density of 2 × 10^4^ cells per well. After 18 to 24 hours, the cells, at 75-90% confluency, were transfected with 200 ng psDARPin-expressing plasmids using lipofectamine 3000 (Invitrogen), following the manufacturer’s instructions.

### Fluorescence microscopy and image analysis

Fluorescence microscopy was performed with a confocal microscope (Carl Zeiss, LSM 980) using a Plan-Apochromat 63x/1.4 Oil DIC M27 objective and a GaAsP-PMT detector. The fluorescent probes were image using pixel dwell times of 0.42 μs and the following settings: mTurq2 (laser excitation: 0.6 mW 445 nm, detection range: 450–485 nm), pdDronpa (laser excitation: 0.03 mW 488 nm, detection range: 500–566 nm), mScarlet (laser excitation: 0.02 mW 561 nm, detection range: 565–758 nm), and Alexa Fluor 647 (laser excitation: 0.41 mW 639 nm, detection range: 646–758 nm). When testing mTurq2 C/N ratios in the basal state. Brief one-frame 488-nm excitation of pdDronpa immediately preceded 445-nm excitation to image mTurq2. Live cell imaging was conducted in an incubation chamber at 37 °C under 5 % CO_2_ atmosphere. Cells for data analysis were randomly selected from three independent experiments with varied protein expression levels. Zen 2.5 (Carl Zeiss) and ImageJ (National Institutes of Health) software were used to analyze mean fluorescence intensities of pdDronpa, mTurq2, and mScarlet following manual segmentation of the nucleus, cytosol, mitochondria, and plasma membrane. Based on these results, the fluorescence ratios including C/N Turq2 ratios for ps3G124 variants, C/N mScarlet ratios for psE40/K27 variants, were calculated as specified in the figures on a per-cell basis. The background signal was determined as the mean fluorescence from a non-cell region of the image and subtracted from all signals prior to calculating the ratio.

### Photoswitching of pdDronpa

To photoswitch pdDronpa from the ON to the OFF state an ROI was drawn around the cells within the field of view and the 488-nm confocal illumination was scanned across the ROI for 10 s using the power listed in the figure captions. Similarly, the reverse OFF to ON photoswitching was performed using 0.56 mW 405-nm laser excitation scanned for 5 s across the ROI.

### EGF-stimulation and late-stage ERK signaling in serum starved HeLa cells

Plasmids encoding ps3G124.ex17 or psK27.ex17 were transfected at 200 ng in Optimem (Gibco) for 4 hours, followed by replacement with DMEM supplemented with 10% FBS for 16–20 hours. 4 hours before cell imaging, the cells were shifted to a FBS-starved medium: two consecutive medium replacements were conducted with FBS-free DMEM, followed by incubation in DMEM containing 0.1% FBS for 4 hours. Images were recorded at 5-minute intervals before and after treatment with 10 ng/mL of Alexa Fluor™ 647-labelled EGF (EGF-AF647, Thermo Scientific) or human EGF (Abcam).

### Statistics and reproducibility

Statistical information is provided in the figure legends. All quantifications are shown as mean ± SD.

## Data availability

The crystal structures of 3G124.ex13 and Off7.ex17 have been deposited in the PDB database under accession codes 8U34 and 8U35, respectively.

## Funding

The project is funded by NIH NIGMS award 1R21GM132687 (MZL), grant NNF18OC0031816 from the Novo Nordisk Foundation and the Stanford Bio-X Program (MW), grant NRF-2021R1C1C2009359 from the National Research Foundation of Korea supported by the Korean MSIT (DS), and HFSP Long-term fellowship LT0062/2022-L (DS). Use of the Stanford Synchrotron Radiation Lightsource, SLAC National Accelerator Laboratory, is supported by the U.S. Department of Energy, Office of Science, Office of Basic Energy Sciences under Contract No. DE-AC02-76SF00515. The SSRL Structural Molecular Biology Program is supported by the DOE Office of Biological and Environmental Research, and by grant P30GM133894 from NIH NIGMS.

### Acknowledgements

We thank Gordon Wang of the Stanford Neuroscience Microscopy Service for valuable advice and Hokyung Kay Chung for generating the U87E stable line while in the Lin laboratory. Part of the work was performed at the Neuroscience Microscopy Facility of the Wu Tsai Neuroscience Institute at Stanford. A portion of this work was performed at the Sarafan ChEM-H Macromolecular Structure Knowledge Center at Stanford. Some of the computing for this project was performed on the Sherlock cluster of the Stanford Research Computing Center. We thank Stanford University for resources and support from all three core facilities.

ERK-KTR was a gift from the Covert lab at Stanford University. The pET His6 TEV LIC cloning vector (2B-T) was a gift from Scott Gradia (Addgene plasmid # 29666 ; http://n2t.net/addgene:29666 ; RRID:Addgene_29666).

## Author contributions

Initial conceptualization of the psDARPin concept was performed by V.D. and M.Z.L, and subsequently refined by M.W. Computational design, expression, and purification of exDARPins were performed by M.W. and V.D. with supervision from P.H. and M.Z.L. exDARPins were crystallized by V.D. with supervision from M.W. Diffraction data were collected and integrated by D.F., and structural models were built by M.W., V.D, and D.F. Protein stability tests were performed by V.D. with training and supervision from M.W. Live-cell experiments of psDARPin function on mTurq2, Erk, and Ras and image analysis were performed by D.S. with supervision from M.W. and M.Z.L. The manuscript was written and graphics were prepared by M.W., D.S., and M.Z.L. All authors reviewed and contributed to the final manuscript.

## Corresponding authors

Correspondence to Michael Westberg and Michael Z. Lin

---

**Extended Data Fig. 1.**
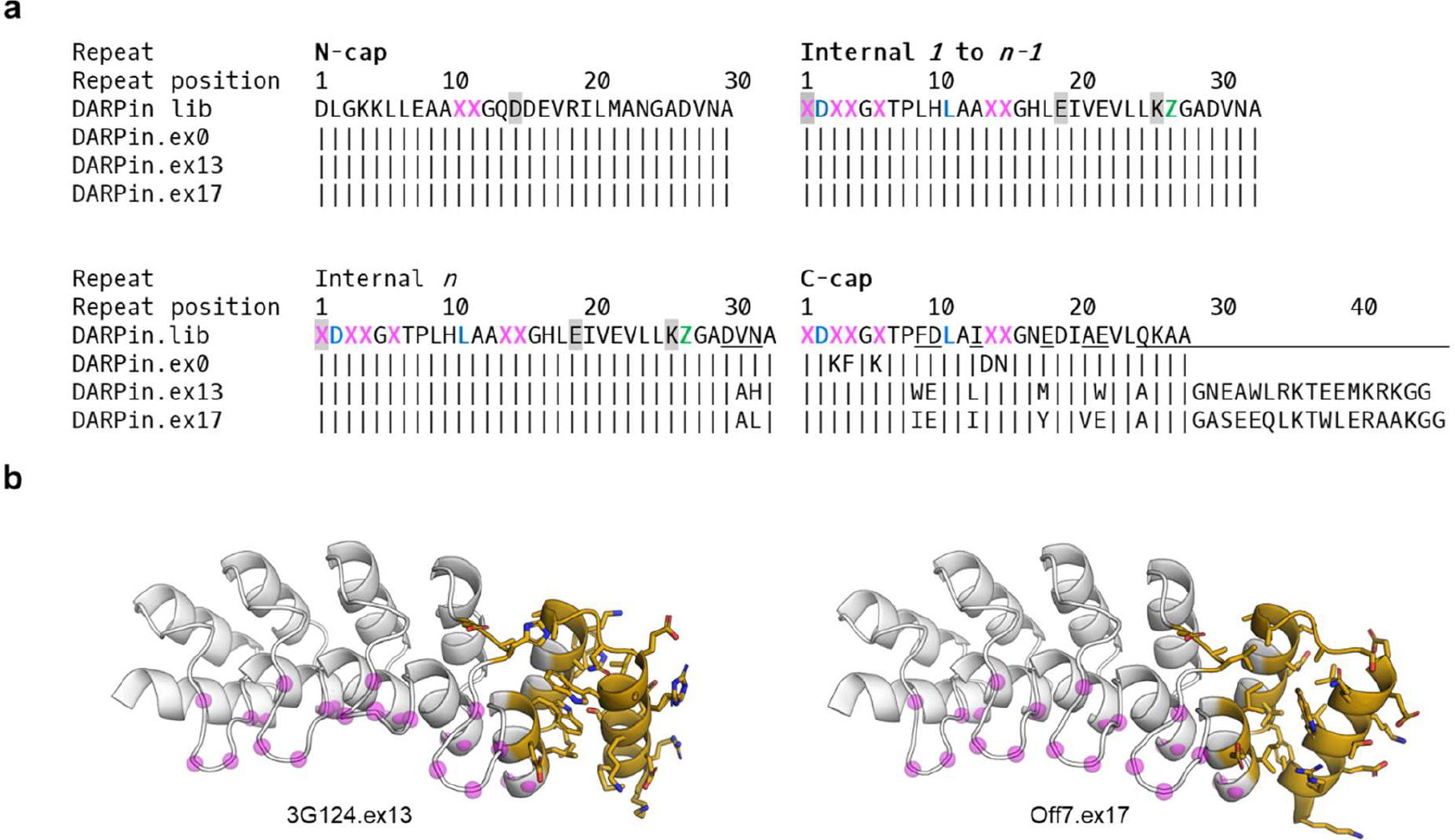
Designing exDARPins while preserving the diversified paratope positions of canonical DARPin libraries. **a**, Sequence alignments comparing the exDARPin designs to a canonical DARPin sequence. DARPin.lib represents an optimized canonical DARPin library sequence that is diversified on the positions marked with X (any amino acid except G/C/P) and Z (any of H/N/Y). The residue positions marked with a magenta X make up the primary paratope positions highlighted in b as magenta spheres. However, random mutagenesis and affinity maturation may lead to DARPins that contain sporadic mutations of conserved residues beyond the initial diversified positions of the library. Over time, several groups have reported optimized C-caps. In this study we used the mut5 cap, here ex0, reported by Interlandi et al.^34^ as a representative optimized canonical C-cap and as a starting point for the computational designs. In the original DARPin libraries^31,82^, no position in the N-cap or C-cap were diversified. However, more recent libraries incorporate diversification of cap positions that structurally correspond to the canonically diversified positions of the internal repeats^14,83,84^. Furthermore, deviating slightly from the canonical paratope pattern^84^, positions 2 and 11 of the internal repeats and C-cap (blue) have also been suggested as sites for library diversification. Thus, we considered all these positions (magenta and blue) as potential paratope residues during our process of computationally designing the extended C-caps of exDARPins. Specifically, only underlined residues were computationally designed with RosettaRemodel. In addition to redesigning the C-cap, we also allowed the residues of positions 30-32 of internal repeat *n* to vary to facilitate interactions that could stabilize the new loop leading into the added helix. Otherwise, all the computationally optimized residues are positioned within, or as an extension of, the C-cap. Sequence alignments of exDARPins DARPin.ex13 and DARPin.ex17 that were used throughout this study show that the extended C-cap designs do not affect any of the library position (magenta and blue). This was the case for all designs. Optimization of the N-cap and internal repeats have previously been reported by mutating the residues highlighted with grey. Particularly, a D15L^85^ mutation in the N-cap as well as E18D^84^ and K25A^84^ mutations in the internal repeats have been reported to optimize DARPin stability. We did not incorporate any of these mutations in this study, however, as is seen from the sequence alignments, exDARPins are fully compatible with these mutations. **b**, Crystal structures of 3G124.ex13 and Off7.ex17 with the computationally optimized positions highlighted in gold (i.e., corresponding to residues underlined in panel a) and library position marked with magenta spheres (i.e., corresponding to residues marked with X in panel a).

**Extended Data Fig. 2.**
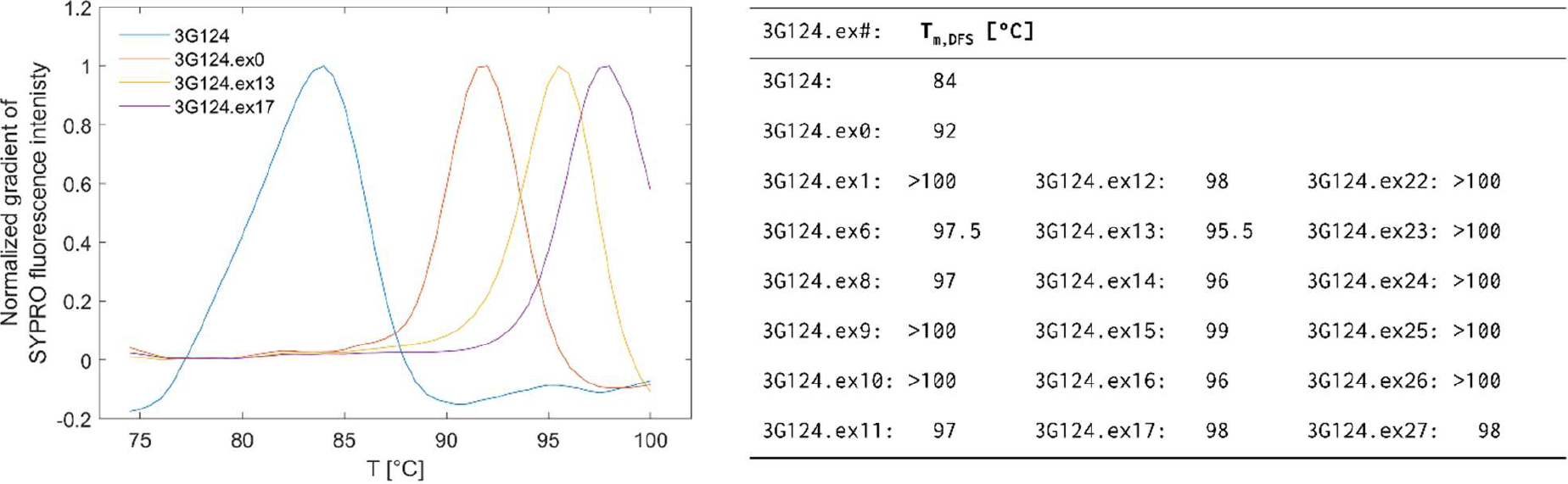
Thermal stability of exDARPins by differential scanning fluorimetry. The thermal stabilities of DARPins were measured using differential scanning fluorimetry. The T_m_ values were determined as the peak position of the gradient of the SYPRO Orange fluorescence signal. 3G124 is the original version of the anti-GFP DARPin reported in Brauchle et al.^32^. that does not contain the optimized C-cap (i.e., ex0) reported by Interlandi et al.^34^. The more stable 3G124.ex0 served as the reference point throughout this study. The computationally designed exDARPins are listed as ex1−ex27.

**Extended Data Fig. 3.**
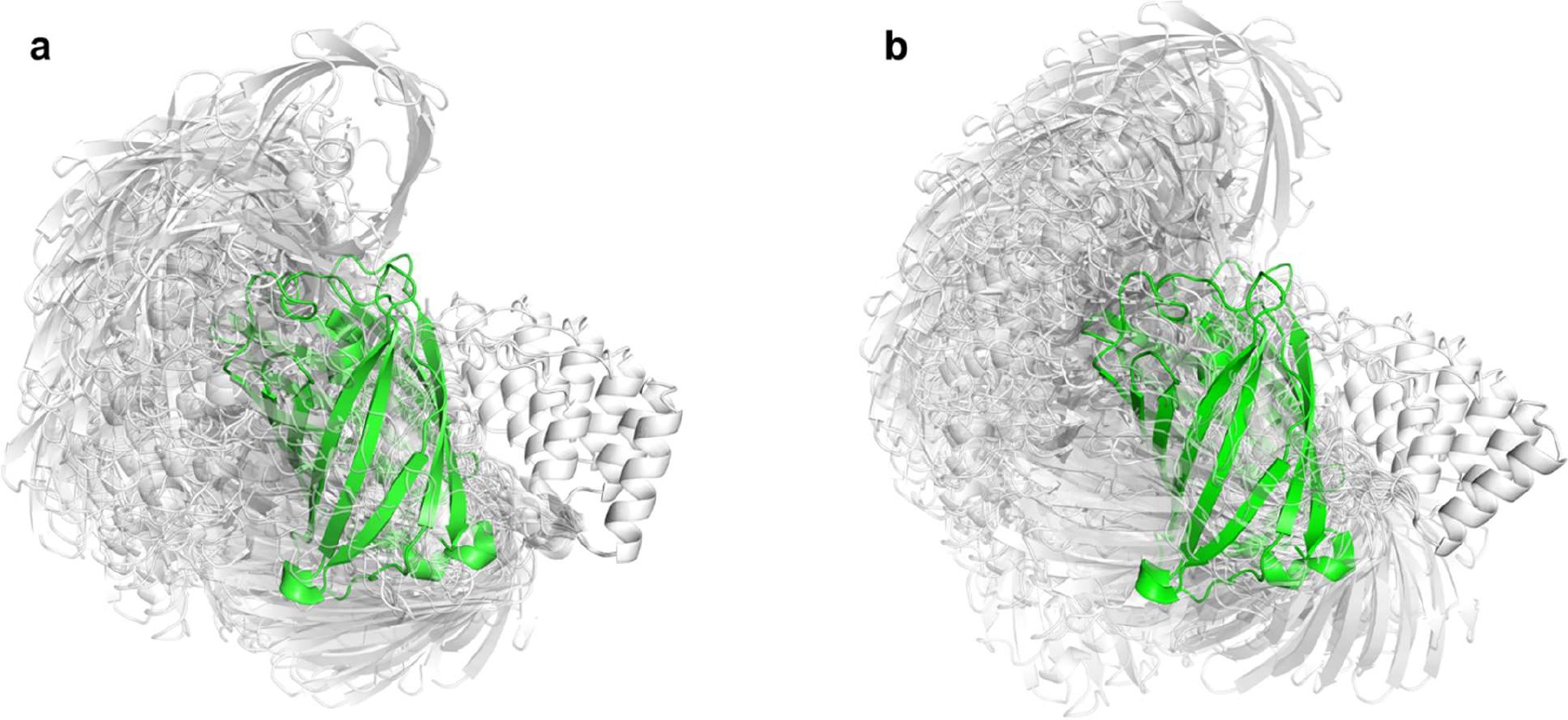
Computational linker screen. RosettaRemodel was used to screen linker lengths by sampling the fusion between pdDronpa1.2 and exDARPins using a chain-closure algorithm^36^. Output structures from the modelling of the 7,5 linker combination, that was used for the majority of the experiments, is shown for **a**, ps3G124.ex13 and **b**, ps3G124.ex17. The psDARPins are shown in transparent white with the new C-terminal in the foreground, and the GFP target is shown in green. For each linker combination, 100 independent trajectories were run, and only the structures of trajectories with successful chain-closure are shown. Based on these simulations, we prioritized linker combinations for which the pdDronpa1.2 cage is centered in front of the exDARPins paratope and with either of the pdDronpa1.2 domains occluding target binding while avoiding strained linkers or clashes.

**Extended Data Fig. 4.**
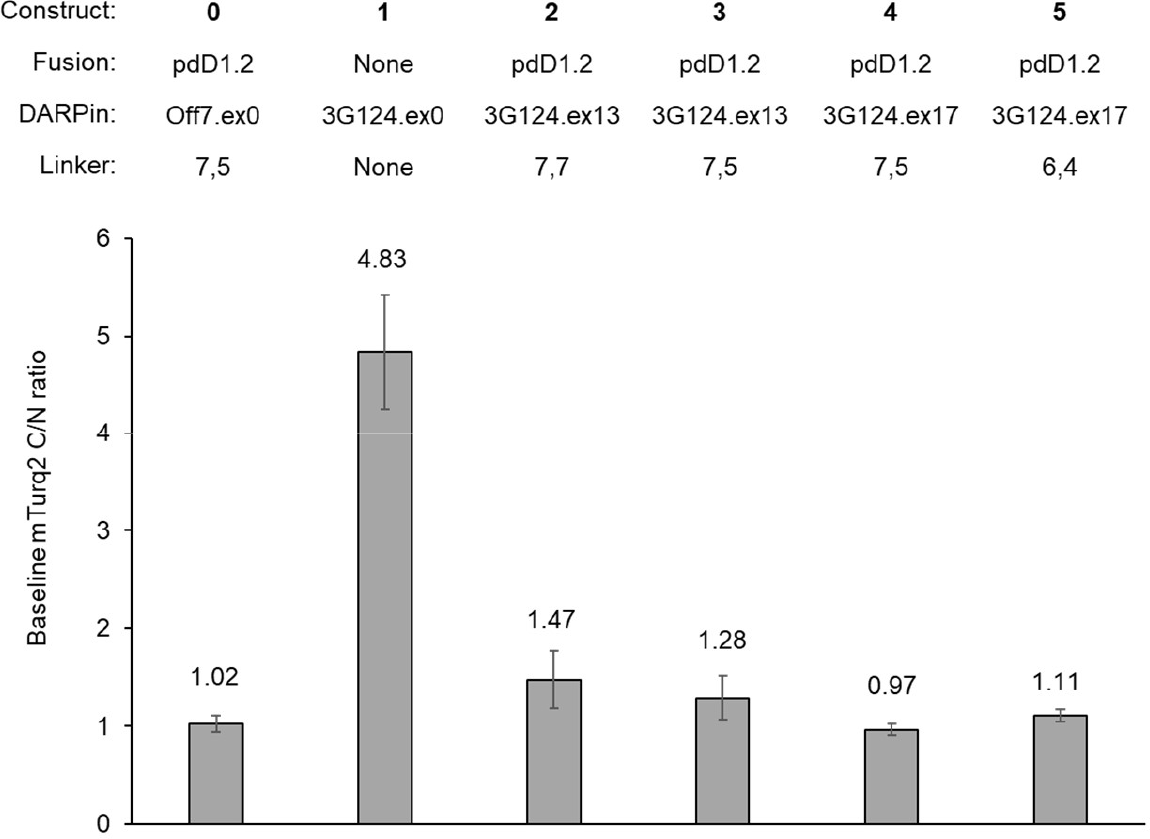
Experimental linker screen. Focused linker screen of caging of ps3G124.ex13 and ps3G124.ex17 under dark basal conditions. NES-fused constructs were co-expressed with mTurq2, and mTurq2-binding was assessed by the mTurq2 C/N ratio at baseline in the absence of illumination. The tested constructs were: **0** (also found in Fig. 2b-c), the extension-free Off7 (Off7.ex0) DARPin recognizing E. coli maltose-binding protein with terminal pdDronpa1.2 (pdD1.2) fusions, as a negative control not expected to bind mTurq2; **1**, (also found in Fig. 2b-c), the extension-free GFP-binding 3G124 (3G124.ex0) without pdD1.2 attachments, as a DARPin expected to constitutively sequester mTurq2 in the cytosol; **2**, 3G124 with designed extension ex13 and terminal pdD1.2 attachments via linkers of 7,7 residues, to test the caging efficiency; **3**, (also found in Fig. 2b-c), 3G124 with designed extension ex13 and terminal pdD1.2 attachments via linkers of 7,5 residues (ps3G124.ex13), to test the caging efficiency; **4**, (also found in Fig. 2b-c), 3G124 with designed extension ex17 and terminal pdD1.2 attachments via linkers of 7,5 residues (ps3G124.ex17), to test the caging efficiency; **5**, 3G124 with designed extension ex13 and terminal pdD1.2 attachments via linkers of 6,4 residues, to test the caging efficiency. For both 3G124.ex13 and 3G124.ex17 the 7,5 linker was the best performing linker combination.

**Extended Data Fig. 5.**
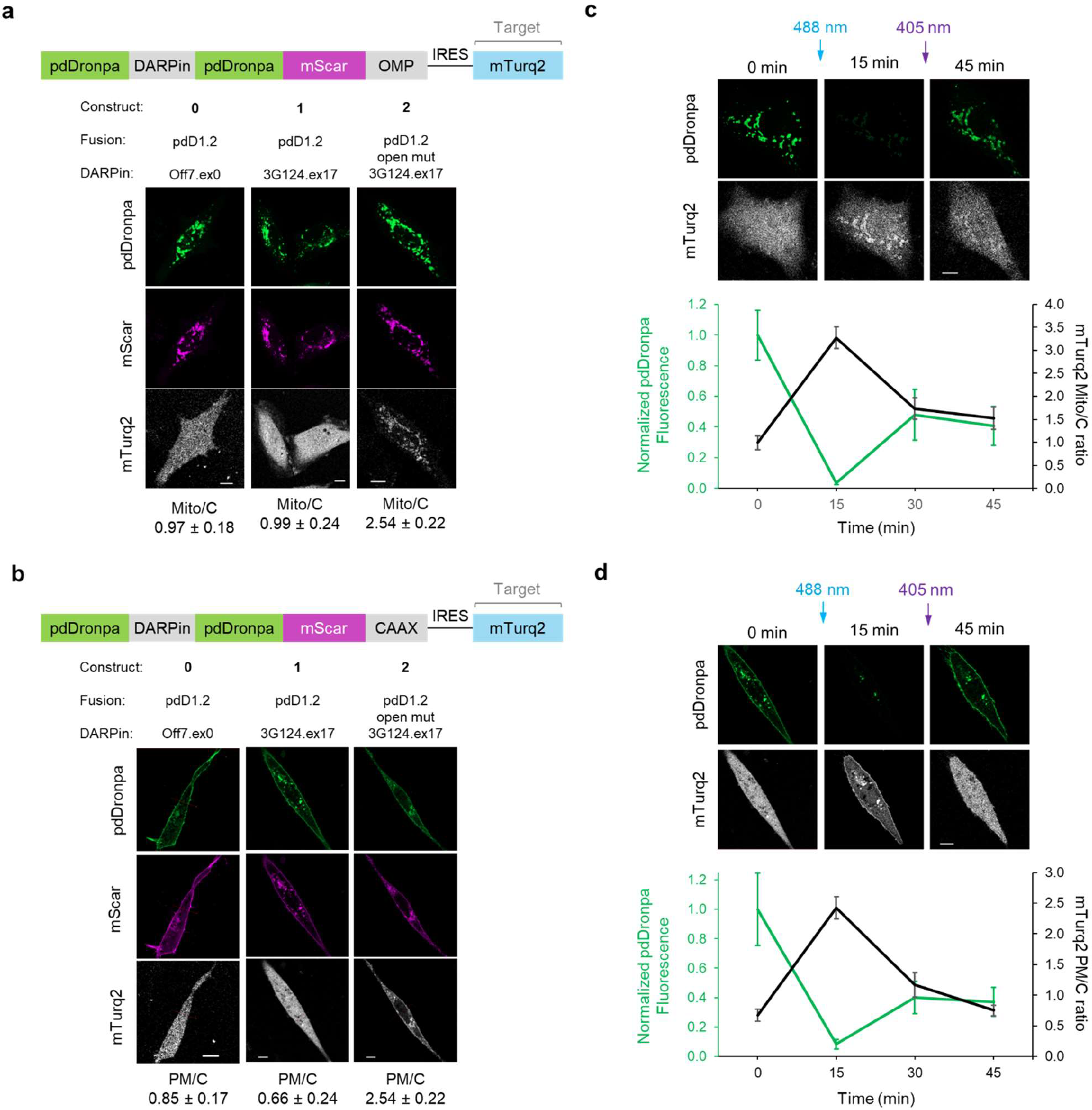
Relocalization of target proteins by subcellularly localized psDARPin. Schematic diagrams of **a**, mitochondria (mito) or **b**, plasma membrane (PM) anchored ps3G124.ex17 constructs and representative images of HeLa cells expressing the various test constructs in the dark. The DARPin fusions localized as expected, and the fusion to pdD1.2 ensure caging of the paratope. Recruitment of mTurq2 to the organelles was quantified as the mTurq2 fluorescence intensity at the mitochondria vs. the cytosol (Mito/C) or the plasma membrane vs. the cytosol (PM/C) (*n,N* = 18 independent cells, 3 independent experiments per construct). Representative images for bidirectional photoswitching of both **c**, mito- and **d**, PM-anchored versions of construct 1 and quantification of the associated fluorescence ratios (*n,N* = 18 independent cells, 3 independent experiments per construct). Following 488-nm illumination (10 s, 0.21 mW), mTurq2 was recruited to the mitochondria or plasma membrane, respectively. The psDARPins were recaged after 405-nm illumination (5 s, 0.56 mW) as seen from the release of mTurq2 back into the cytosol. In all instances, scale bars are 10 μm, mean ratios are reported, and error bars are standard deviation (SD) calculated using *n*.

**Extended Data Fig. 6.**
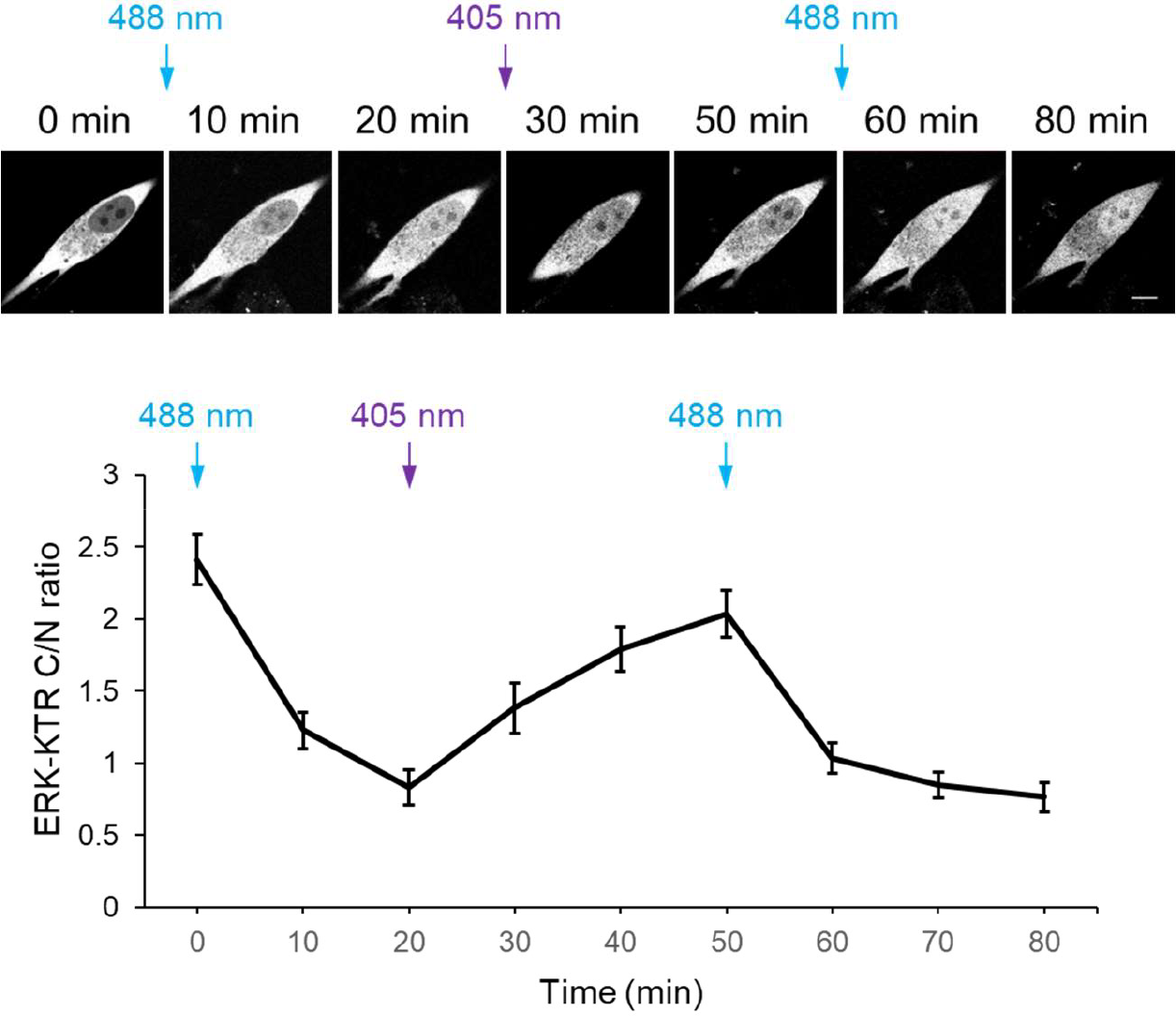
Photoinduced inhibition of endogenous ERK and Ras with psDARPins. Top, representative mScarlet images from U87E cells expressing psE40.ex13 that was sequentially switched with 488-nm light (10 s, 0.21 mW) and 405-nm light (5 s, 0.56 mW). Bottom, quantification of the associated ERK-KTR C/N ratios (*n,N* = 18 independent cells, 3 independent experiments per construct). In all instances, scale bars are 10 μm, mean C/N ratios are reported, and error bars are standard deviation (SD) calculated using *n*.

## References

1. Wang, H., La Russa, M. & Qi, L. S. CRISPR/Cas9 in Genome Editing and Beyond. Annu. Rev. Biochem. 85, 227–264 (2016).

2. Dykxhoorn, D. M. & Lieberman, J. The Silent Revolution: RNA Interference as Basic Biology, Research Tool, and Therapeutic. Annu. Rev. Med. 56, 401–423 (2005).

3. Yu, X., Yang, Y.-P., Dikici, E., Deo, S. K. & Daunert, S. Beyond Antibodies as Binding Partners: The Role of Antibody Mimetics in Bioanalysis. Annu. Rev. Anal. Chem. 10, 293–320 (2017).

4. Hosse, R. J. A new generation of protein display scaffolds for molecular recognition. Protein Sci. 15, 14–27 (2006).

5. Sha, F., Salzman, G., Gupta, A. & Koide, S. Monobodies and other synthetic binding proteins for expanding protein science: Monobodies and Other Synthetic Binding Proteins. Protein Sci. 26, 910–924 (2017).

6. Mohan, K. et al. Topological control of cytokine receptor signaling induces differential effects in hematopoiesis. Science 364, eaav7532 (2019).

7. Steiner, D., Forrer, P., Stumpp, M. T. & Plückthun, A. Signal sequences directing cotranslational translocation expand the range of proteins amenable to phage display. Nat. Biotechnol. 24, 823–831 (2006).

8. Dong, J.-X. et al. A toolbox of nanobodies developed and validated for use as intrabodies and nanoscale immunolabels in mammalian brain neurons. eLife 8, e48750 (2019).

9. Stern, L. A., Case, B. A. & Hackel, B. J. Alternative non-antibody protein scaffolds for molecular imaging of cancer. Curr. Opin. Chem. Eng. 2, 425–432 (2013).

10. Gebauer, M. & Skerra, A. Engineered Protein Scaffolds as Next-Generation Therapeutics. Annu. Rev. Pharmacol. Toxicol. 60, 391–415 (2020).

11. Wheeler, Y. Y., Chen, S.-Y. & Sane, D. C. Intrabody and intrakine strategies for molecular therapy. Mol. Ther. 8, 355–366 (2003).

12. Sha, F., Salzman, G., Gupta, A. & Koide, S. Monobodies and other synthetic binding proteins for expanding protein science: Monobodies and Other Synthetic Binding Proteins. Protein Sci. 26, 910–924 (2017).

13. Marschall, A. L., Dübel, S. & Böldicke, T. Specific in vivo knockdown of protein function by intrabodies. mAbs 7, 1010–1035 (2015).

14. Plückthun, A. Designed Ankyrin Repeat Proteins (DARPins): Binding Proteins for Research, Diagnostics, and Therapy. Annu. Rev. Pharmacol. Toxicol. 55, 489–511 (2015).

15. Wild, P., Eapen, R., Forrer, P. & Jost, C. The DARPin Encyclopedia: from Basic Research towards Therapeutics. preprint.org (2022) doi:10.20944/preprints202206.0147.v3.

16. Lucchi, R., Bentanachs, J. & Oller-Salvia, B. The Masking Game: Design of Activatable Antibodies and Mimetics for Selective Therapeutics and Cell Control. ACS Cent. Sci. 7, 724–738 (2021).

17. Yu, D. et al. Optogenetic activation of intracellular antibodies for direct modulation of endogenous proteins. Nat. Methods (2019) doi:10.1038/s41592-019-0592-7.

18. Gil, A. A. et al. Optogenetic control of protein binding using light-switchable nanobodies. Nat. Commun. 11, 4044 (2020).

19. He, L., Tan, P., Huang, Y. & Zhou, Y. Design of Smart Antibody Mimetics with Photosensitive Switches. Adv. Biol. 5, 2000541 (2021).

20. Woloschuk, R. M. et al. Structure-based design of a photoswitchable affibody scaffold. Protein Sci. (2021) doi:10.1002/pro.4196.

21. Brechun, K. E. et al. Detection of Incorporation of p -Coumaric Acid into Photoactive Yellow Protein Variants in Vivo. Biochemistry 58, 2682–2694 (2019).

22. Ryu, H. et al. Combinatorial effects of RhoA and Cdc42 on the actin cytoskeleton revealed by photoswitchable GEFs. Sens. Actuators B Chem. 369, 132316 (2022).

23. Zhou, X. X., Chung, H. K., Lam, A. J. & Lin, M. Z. Optical Control of Protein Activity by Fluorescent Protein Domains. Science 338, 810–814 (2012).

24. Zhou, X. X., Fan, L. Z., Li, P., Shen, K. & Lin, M. Z. Optical control of cell signaling by single-chain photoswitchable kinases. Science 355, 836–842 (2017).

25. Zhou, X. X. et al. A Single-Chain Photoswitchable CRISPR-Cas9 Architecture for Light-Inducible Gene Editing and Transcription. ACS Chem. Biol. 13, 443–448 (2018).

26. Ju, J. et al. Optical regulation of endogenous RhoA reveals selection of cellular responses by signal amplitude. Cell Rep. 40, 111080 (2022).

27. Patel, A. L. et al. Optimizing photoswitchable MEK. Proc. Natl. Acad. Sci. 116, 25756–25763 (2019).

28. Jones, T., Liu, A. & Cui, B. Light-Inducible Generation of Membrane Curvature in Live Cells with Engineered BAR Domain Proteins. ACS Synth. Biol. 9, 893–901 (2020).

29. Boersma, Y. L. Advances in the Application of Designed Ankyrin Repeat Proteins (DARPins) as Research Tools and Protein Therapeutics. in Protein Scaffolds (ed. Udit, A. K.) vol. 1798 307–327 (Springer New York, 2018).

30. Forrer, P., Stumpp, M. T., Binz, H. K. & Plückthun, A. A novel strategy to design binding molecules harnessing the modular nature of repeat proteins. FEBS Lett. 539, 2–6 (2003).

31. Binz, H. K. et al. High-affinity binders selected from designed ankyrin repeat protein libraries. Nat. Biotechnol. 22, 575–582 (2004).

32. Brauchle, M. et al. Protein interference applications in cellular and developmental biology using DARPins that recognize GFP and mCherry. Biol. Open 3, 1252–1261 (2014).

33. Huang, P.-S. et al. RosettaRemodel: A Generalized Framework for Flexible Backbone Protein Design. PLoS ONE 6, e24109 (2011).

34. Interlandi, G., Wetzel, S. K., Settanni, G., Plückthun, A. & Caflisch, A. Characterization and Further Stabilization of Designed Ankyrin Repeat Proteins by Combining Molecular Dynamics Simulations and Experiments. J. Mol. Biol. 375, 837–854 (2008).

35. Kramer, M. A., Wetzel, S. K., Plückthun, A., Mittl, P. R. E. & Grütter, M. G. Structural determinants for improved stability of designed ankyrin repeat proteins with a redesigned C-Capping Module. J. Mol. Biol. 404, 381–391 (2010).

36. Rose, J. C. et al. A computationally engineered RAS rheostat reveals RAS–ERK signaling dynamics. Nat. Chem. Biol. 13, 119–126 (2017).

37. Goedhart, J. et al. Structure-guided evolution of cyan fluorescent proteins towards a quantum yield of 93%. Nat. Commun. 3, 751 (2012).

38. Hansen, S. et al. Design and applications of a clamp for Green Fluorescent Protein with picomolar affinity. Sci. Rep. 7, 1–16 (2017).

39. Lavoie, H., Gagnon, J. & Therrien, M. ERK signalling: a master regulator of cell behaviour, life and fate. Nat. Rev. Mol. Cell Biol. 21, 607–632 (2020).

40. Punekar, S. R., Velcheti, V., Neel, B. G. & Wong, K.-K. The current state of the art and future trends in RAS-targeted cancer therapies. Nat. Rev. Clin. Oncol. 19, 637–655 (2022).

41. Samatar, A. A. & Poulikakos, P. I. Targeting RAS–ERK signalling in cancer: promises and challenges. Nat. Rev. Drug Discov. 13, 928–942 (2014).

42. Rauen, K. A. The RASopathies. Annu. Rev. Genomics Hum. Genet. 14, 355–369 (2013).

43. Kummer, L. et al. Structural and functional analysis of phosphorylation-specific binders of the kinase ERK from designed ankyrin repeat protein libraries. Proc. Natl. Acad. Sci. 109, (2012).

44. Stephens, E. A. et al. Engineering Single Pan-Specific Ubiquibodies for Targeted Degradation of All Forms of Endogenous ERK Protein Kinase. ACS Synth. Biol. 10, 2396–2408 (2021).

45. Regot, S., Hughey, J. J., Bajar, B. T., Carrasco, S. & Covert, M. W. High-sensitivity measurements of multiple kinase activities in live single cells. Cell 157, 1724–1734 (2014).

46. Moore, A. R., Rosenberg, S. C., McCormick, F. & Malek, S. RAS-targeted therapies: is the undruggable drugged? Nat. Rev. Drug Discov. 19, 533–552 (2020).

47. Welsch, M. E. et al. Multivalent Small-Molecule Pan-RAS Inhibitors. Cell 168, 878–889.e29 (2017).

48. Kessler, D. et al. Drugging an undruggable pocket on KRAS. Proc. Natl. Acad. Sci. 116, 15823–15829 (2019).

49. Coley, A. B. et al. Pan-RAS inhibitors: Hitting multiple RAS isozymes with one stone. in Advances in Cancer Research vol. 153 131–168 (Elsevier, 2022).

50. Spencer-Smith, R. et al. Inhibition of RAS function through targeting an allosteric regulatory site. Nat. Chem. Biol. 13, 62–68 (2017).

51. Guillard, S. et al. Structural and functional characterization of a DARPin which inhibits Ras nucleotide exchange. Nat. Commun. 8, 1–11 (2017).

52. Khan, I. et al. Identification of the nucleotide-free state as a therapeutic vulnerability for inhibition of selected oncogenic RAS mutants. Cell Rep. 38, 110322 (2022).

53. Wallon, L. et al. Inhibition of RAS-driven signaling and tumorigenesis with a pan-RAS monobody targeting the Switch I/II pocket. Proc. Natl. Acad. Sci. 119, e2204481119 (2022).

54. Bery, N. et al. KRAS-specific inhibition using a DARPin binding to a site in the allosteric lobe. Nat. Commun. 10, 0–10 (2019).

55. Surve, S., Watkins, S. C. & Sorkin, A. EGFR-RAS-MAPK signaling is confined to the plasma membrane and associated endorecycling protrusions. J. Cell Biol. 220, e202107103 (2021).

56. Pinilla-Macua, I., Watkins, S. C. & Sorkin, A. Endocytosis separates EGF receptors from endogenous fluorescently labeled HRas and diminishes receptor signaling to MAP kinases in endosomes. Proc. Natl. Acad. Sci. 113, 2122–2127 (2016).

57. Sasagawa, S., Ozaki, Y., Fujita, K. & Kuroda, S. Prediction and validation of the distinct dynamics of transient and sustained ERK activation. Nat. Cell Biol. 7, 365–373 (2005).

58. Weeks, R., Zhou, X., Yuan, T. L. & Zhang, J. Fluorescent Biosensor for Measuring Ras Activity in Living Cells. J. Am. Chem. Soc. 144, 17432–17440 (2022).

59. Zhang, J. Z. et al. Computationally designed sensors for endogenous Ras activity reveal signaling effectors within oncogenic granules. biorxiv (2023) doi:10.1101/2022.11.22.517009.

60. Bivona, T. G., Quatela, S. & Philips, M. R. Analysis of Ras Activation in Living Cells with GFP-RBD. in Methods in Enzymology vol. 407 128–143 (Elsevier, 2006).

61. Dessauges, C. et al. Optogenetic actuator – ERK biosensor circuits identify MAPK network nodes that shape ERK dynamics. Mol. Syst. Biol. 18, (2022).

62. Toettcher, J. E., Weiner, O. D. & Lim, W. A. Using optogenetics to interrogate the dynamic control of signal transmission by the Ras/Erk module. Cell 155, 1422–1434 (2013).

63. Komatsu, T. et al. Organelle-specific, rapid induction of molecular activities and membrane tethering. Nat. Methods 7, 206–208 (2010).

64. Maryu, G., Matsuda, M. & Aoki, K. Multiplexed Fluorescence Imaging of ERK and Akt Activities and Cell-cycle Progression. Cell Struct. Funct. 41, 81–92 (2016).

65. Redchuk, T. A. et al. Optogenetic regulation of endogenous proteins. Nat. Commun. 11, (2020).

66. Carrasco-López, C. et al. Development of light-responsive protein binding in the monobody non-immunoglobulin scaffold. Nat. Commun. 11, 4045 (2020).

67. Mitchell, L. S. & Colwell, L. J. Comparative analysis of nanobody sequence and structure data. Proteins Struct. Funct. Bioinforma. 86, 697–706 (2018).

68. Fu, L., Chen, S., He, G., Chen, Y. & Liu, B. Targeting Extracellular Signal-Regulated Protein Kinase 1/2 (ERK1/2) in Cancer: An Update on Pharmacological Small-Molecule Inhibitors. J. Med. Chem. 65, 13561–13573 (2022).

69. Bugaj, L. J. et al. Cancer mutations and targeted drugs can disrupt dynamic signal encoding by the Ras-Erk pathway. Science 361, eaao3048 (2018).

70. McFann, S. E., Shvartsman, S. Y. & Toettcher, J. E. Putting in the Erk: Growth factor signaling and mesoderm morphogenesis. in Current Topics in Developmental Biology vol. 149 263–310 (Elsevier, 2022).

71. Meksiriporn, B. et al. A survival selection strategy for engineering synthetic binding proteins that specifically recognize post-translationally phosphorylated proteins. Nat. Commun. 10, (2019).

72. Liu, X. & Ciulli, A. Proximity-Based Modalities for Biology and Medicine. ACS Cent. Sci. 9, 1269–1284 (2023).

73. Kabsch, W. XDS. Acta Crystallogr. D Biol. Crystallogr. 66, 125–132 (2010).

74. Winter, G. et al. DIALS: Implementation and evaluation of a new integration package. Acta Crystallogr. Sect. Struct. Biol. 74, 85–97 (2018).

75. Evans, P. R. & Murshudov, G. N. How good are my data and what is the resolution? Acta Crystallogr. D Biol. Crystallogr. 69, 1204–1214 (2013).

76. Collaborative Computational Project. The CCP4 suite: Programs for protein crystallography. Acta Crystallogr. D Biol. Crystallogr. 50, 760–763 (1994).

77. Potterton, L. et al. CCP4i2: the new graphical user interface to the CCP4 program suite. Acta Crystallogr. Sect. Struct. Biol. 74, 68–84 (2018).

78. McCoy, A. J. et al. Phaser crystallographic software. J. Appl. Crystallogr. 40, 658–674 (2007).

79. Murshudov, G. N., Vagin, A. A. & Dodson, E. J. Refinement of macromolecular structures by the maximum-likelihood method. Acta Crystallogr. D Biol. Crystallogr. 53, 240–255 (1997).

80. Emsley, P. & Cowtan, K. Coot: Model-building tools for molecular graphics. Acta Crystallogr. D Biol. Crystallogr. 60, 2126–2132 (2004).

81. Guo, G. et al. Ligand-Independent EGFR Signaling. Cancer Res. 75, 3436–3441 (2015).

82. Binz, H. K., Stumpp, M. T., Forrer, P., Amstutz, P. & Plückthun, A. Designing repeat proteins: Well-expressed, soluble and stable proteins from combinatorial libraries of consensus ankyrin repeat proteins. J. Mol. Biol. 332, 489–503 (2003).

83. Schilling, J., Schöppe, J. & Plückthun, A. From DARPins to LoopDARPins: Novel LoopDARPin design allows the selection of low picomolar binders in a single round of ribosome display. J. Mol. Biol. 426, 691–721 (2014).

84. Seeger, M. A. et al. Design, construction, and characterization of a second-generation DARPin library with reduced hydrophobicity: Second-Generation DARPin Library. Protein Sci. 22, 1239–1257 (2013).

85. Schilling, J. et al. Thermostable designed ankyrin repeat proteins (DARPins) as building blocks for innovative drugs. J. Biol. Chem. 298, 1–12 (2022).

